# Differential gene expression analysis tools exhibit substandard performance for long non-coding RNA–sequencing data

**DOI:** 10.1101/220129

**Authors:** Alemu Takele Assefa, Katrijn De Paepe, Celine Everaert, Pieter Mestdagh, Olivier Thas, Jo Vandesompele

**Author notes:** these authors contributed equally to this work (Alemu Takele Assefa, Olivier Thas, Jo Vandesompele).

## Abstract

**Background:** Protein-coding RNAs (mRNA) have been the primary target of most transcriptome studies in the past, but in recent years, attention has expanded to include long non-coding RNAs (lncRNA). lncRNAs are typically expressed at low levels, and are inherently highly variable. This is a fundamental challenge for differential expression (DE) analysis. In this study, the performance of 14 popular tools for testing DE in RNA-seq data along with their normalization methods is comprehensively evaluated, with a particular focus on lncRNAs and low abundant mRNAs.

**Results:** Thirteen performance metrics were used to evaluate DE tools and normalization methods using simulations and analyses of six diverse RNA-seq datasets. Non-parametric procedures are used to simulate gene expression data in such a way that realistic levels of expression and variability are preserved in the simulated data. Throughout the assessment, we kept track of the results for mRNA and lncRNA separately. All statistical models exhibited inferior performance for lncRNAs compared to mRNAs across all simulated scenarios and analysis of benchmark RNA-seq datasets. No single tool uniformly outperformed the others.

**Conclusion:** Overall, the linear modeling with empirical Bayes moderation (limma) and the nonparametric approach (SAMSeq) showed best performance: good control of the false discovery rate (FDR) and reasonable sensitivity. However, for achieving a sensitivity of at least 50%, more than 80 samples are required when studying expression levels in a realistic clinical settings such as in cancer research. About half of the methods showed severe excess of false discoveries, making these methods unreliable for differential expression analysis and jeopardizing reproducible science. The detailed results of our study can be consulted through a user-friendly web application, http://statapps.ugent.be/tools/AppDGE/

## Introduction

Proteins and messenger RNAs (mRNA) have been the primary target of transcriptome studies. However, RNA sequencing technology has revealed that the human genome is pervasively transcribed, resulting in thousands of novel non-coding RNA genes. Hence, attention is expanding to one of the most poorly understood, yet most common RNA species: long non-coding RNAs (lncRNAs)^1,2^. These lncRNAs form a large and diverse class of transcribed RNA molecules, constituting up to 70% of the transcriptome with a defined length of ≥ 200 nucleotides. While they do not encode for proteins, lncRNAs are strong regulators of gene expression^3^. The discovery and study of lncRNAs is of major relevance to human health and disease because they represent an extensive, largely unexplored, and functional component of the genome^3^–^7^. In contrast to mRNAs, lncRNAs are generally expressed in low amounts, typically an order of magnitude lower than mRNA expression levels^2,5,8–10^. Furthermore, several studies demonstrated that lncRNA expression levels are very noisy, which is a characteristic shared with low count data from massively parallel RNA sequencing^10^–^13^.

Following the advent of RNA-sequencing (RNA-seq) technologies, several statistical tools for differential gene expression (DGE) analysis have been introduced. However, low and noisy read counts, such as those coming from lncRNAs, are potentially challenging the tools^14,15^. For example, it is commonly observed that low count genes or transcripts show large variability of the fold-change estimates and thus exhibit inherently noisier inferential behavior. The majority of the methods suggest removal of low expressed genes before the start of the data analysis, but this procedure essentially blocks the researchers from studying lncRNAs. In our study, no such severe filtering was applied, leaving almost all lncRNAs in the dataset. To our knowledge, there is no statistical method particularly developed for the analysis of lncRNA-seq data and therefore transcriptome studies make use of statistical methods that assume sufficient expression levels. In this paper, we evaluated and compared the performance of many popular statistical methods (see Table 1) developed for testing DGE of RNA-seq data (hereafter referred to as “DE tools” or “DE methods”), with special emphasis on lncRNAs and low abundant mRNAs. All tools considered in this study are popular (in terms of number of citation), available as R software packages^16^, and use gene or transcript level read counts as input. Our conclusions are based on 6 RNA-seq datasets and many realistic simulations, representing various typical gene expression experiments.

**Table 1.** List of DE tools selected for comparison, along with normalization method, brief description, R software package name, and number of citations.

| DE method | normalization method | description | citations<sup>b</sup> | R package (version) | citations-2<sup>c</sup> |
| --- | --- | --- | --- | --- | --- |
| edgeR exact<sup>24</sup> | TMM<sup>25</sup> | Exact test based on Negative Binomial (NB) distribution and empirical Bayes (EB) moderated dispersion estimate | 333 | *edgeR* (3.14.0)<sup>26</sup> | 4526 |
| eddeR GLM<sup>27</sup> | TMM | Generalized linear model (GLM) for NB family and EB moderated dispersion estimate | 442 |  |  |
| edgeR robust<sup>28</sup> | TMM | GLM for NB family with weights to individual observations and EB moderated dispersion estimate | 78 |  |  |
| edgeR QL<sup>29</sup> | TMM | Quasi-likelihood (QL) method with EB moderated QL dispersion estimate | 6 |  |  |
| DESeq<sup>30</sup> | internal<sup>d</sup> | Exact test based on NB distribution with mean-variance modeling to estimate gene specific dispersion parameter | 4,169 | *DESeq* (1.24.0) |  |
| DESeq2<sup>14</sup> | DESeq | GLM for NB family with moderated dispersion and log-fold-change estimates | 1,910 | *DESeq2* (1.12.4) |  |
| limma + QN (limmaQN)<sup>17</sup> | QN<sup>31</sup> | QN was performed on the log2 transformed gene count and linear modeling is applied to test DGE with EB moderated variance estimation | 226 | *limma* (3.28.21)<sup>32</sup> | 1,329 |
| limma + VST (limmaVst)<sup>18</sup> | DESeq | Variance stabilizing transformation (VST) (from *DESeq* package) was applied on the normalized counts and linear modeling is applied to test DGE with EB moderated variance estimation | 257 |  |  |
| limmaVoom<sup>33</sup> | TMM | Calculates weights to each log2 scale normalized counts from observed mean–variance relationship prior to linear modeling | 505 |  |  |
| limmaVoom + Quality Weight (QW)<sup>34</sup> | TMM | Down-weighting observations from more ‘variable’ samples along with observation level weights obtained from the mean–variance relationship and then applies linear modeling | 16 |  |  |
| baySeq<sup>35</sup> | TMM | Posterior likelihoods of DGE using empirical Bayesian methods with prior distributions are estimated empirically | 297 | *baySeq* (2.6.0) |  |
| SAMSeq<sup>23</sup> | internal<sup>d</sup> | A non-parametric method based on Wilcoxon rank sum statistic and a re-sampling procedure of read fragments to calculate sequencing depths | 126 | *samr* (2.0) |  |
| PoissonSeq<sup>36</sup> | internal<sup>d</sup> | Estimates sequencing depths using poisson goodness-of-fit statistic, and then applies poisson log-linear model | 84 | *PoissonSeq* (1.1.2) |  |
| QuasiSeq<sup>a37</sup> | TMM | Tests DGE using Quasi-negative binomial models with ‘QL’, ‘QLShrink’ and ‘QLSpline’ methods. | 44 | *QuasiSeq* (1.0.8) |  |

<sup>a</sup> Based on the recommendation from the original paper, we used ‘NegBinQLSpline’.
<sup>b</sup> As reported by Web of Science (Nov 10, 2017).
<sup>c</sup> If the package has different published documentation. As reported by Web of Science (Nov 10, 2017).
<sup>d</sup> The normalization method is called ‘internal’ if it is not independently reported. We use the name of the tool to refer to the normalization method.
(TMM = trimmed mean of M values, QN = quantile normalization, VST =variance stabilizing transformation)

Previous comparative studies of DE tools^15,17–22^ focused on mRNA and some of these concluded that DE tools show inferior performance for genes or transcripts with low counts. We extend previous studies by including lncRNAs and low expressed mRNAs separately. Our results are based on diverse types of RNA-seq datasets that vary with respect to their biological and technical features, such as species (human, mouse, rat), experimental design (control versus treatment, diseased versus non-diseased, and tissues comparison), and level of biological variability. We assess the degree of concordance among results returned from the DE tools, and we study important statistical properties of DE tools, such as their ability to control the false discovery rate and their sensitivity for the detection of differential expression. The latter are empirically investigated using a nonparametric resampling-based simulation procedure. The simulation method essentially resamples data from a real RNA-seq dataset to create realistic gene expression scenarios. Consequently, our results reflect the genuine behavior of the DE tools under study, in contrast to simulation studies based on parametric assumptions. By starting from a variety of real and representative RNA-seq datasets, the scope of our findings is wide. To our knowledge, our study is the largest empirical evaluation study conducted so far, both in terms of the number of real datasets used and performance metrics evaluated as in the number of DE tools included (Figure A2 of Additional File 1). Note that the evaluation of a method that relies on a parametric assumption (e.g. edgeR, DESeq and DESeq2 assume a negative binomial distribution) by means of simulated counts using the same distribution as used in^18^, will give too optimistic results. Moreover, these results do not reflect a realistic setting because the distributional assumption cannot be expected to hold in general^23^.

Our study consists of three paths: first we evaluated various normalization procedures, second we compared the performance of DE tools across various publicly available RNA-seq datasets, and third we used simulation procedures to evaluate and compare the performance of the tools under a variety of gene expression experiment scenarios (Figure 1), like sample size and fraction of DE genes. We applied most DE tools without modifying their default settings, except for few tools for which minor modifications were made to allow them to work for low abundant genes such as lncRNAs (see Methods section for details).

**Figure 1.**
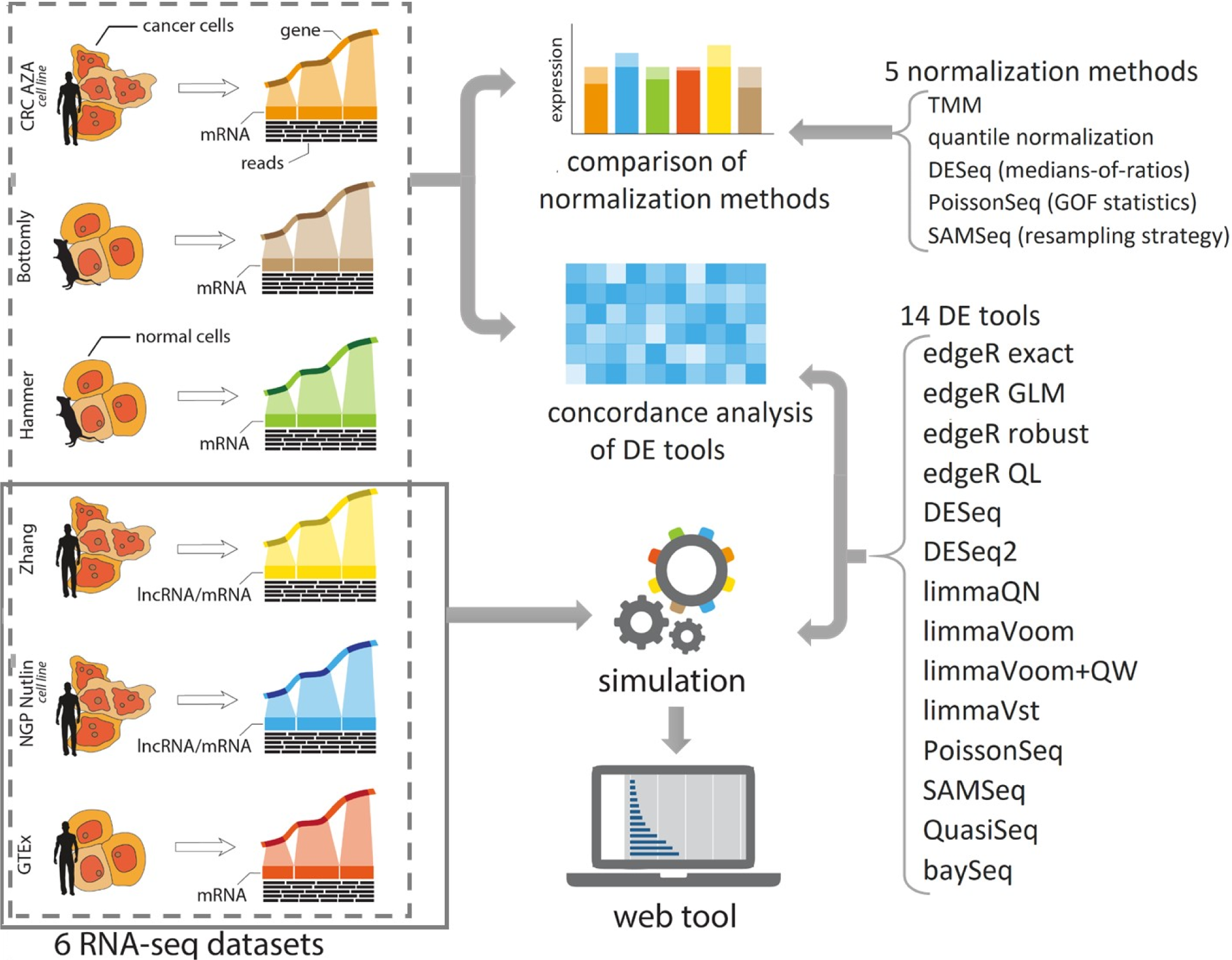
DE tool assessment work flow. The study has three components: evaluation of five normalization methods, concordance analysis of 14 DE tools, and a simulation study to explore statistical properties of 13 DE tools, such as their ability to control the false discovery rate and their sensitivity for the detection of differential expression. Six diverse benchmark RNA-seq datasets are used for comparison of normalization methods and concordance analysis of DE tools. Data are obtained from two cultured cell line datasets (CRC AZA and NGP Nutlin), inbred animals (Bottomly and Hammer), normal human tissues (GTEx), and human cancer cells (Zhang). The simulation study uses three source datasets: GTEx, Zhang, and NGP Nutlin. Results of the simulation study are made available through a user-friendly web application.

## Results and discussion

### RNA-seq datasets

Six publicly available benchmark RNA-seq datasets were used for the concordance analysis. Three of them were used as source datasets for generating non-parametric simulated data. The description of the datasets can be found in the Methods section; a summary is presented in Table 2.

**Table 2.**
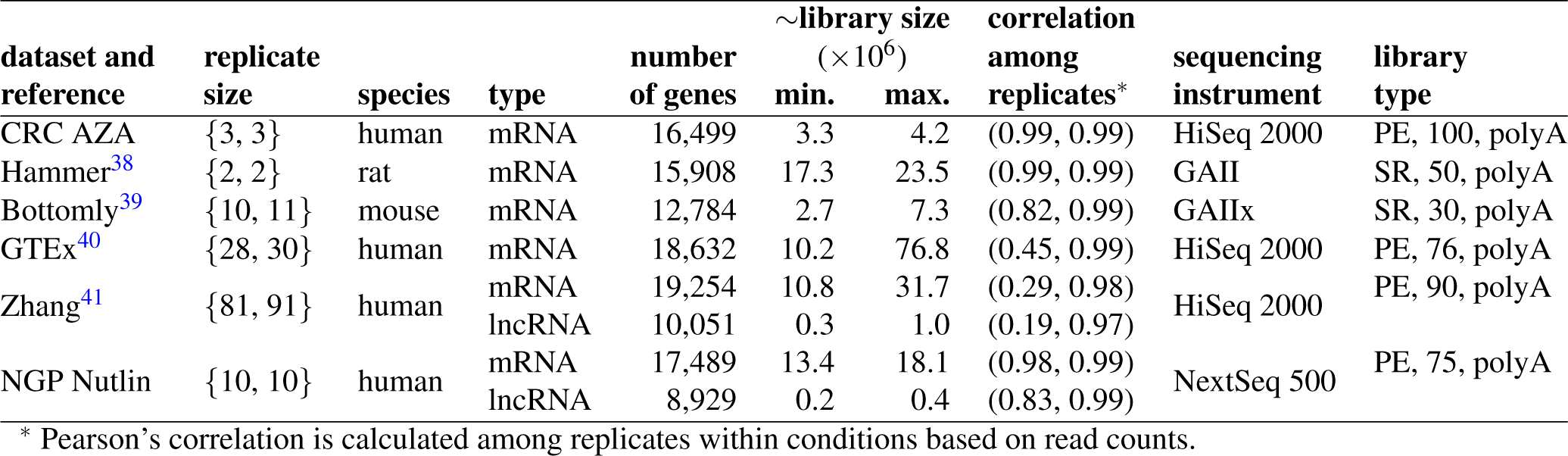
Summary of datasets including the species, gene biotype, number of genes (annotated genes with at least 1 count in each condition), number of replicates per condition, library size (minimum, maximum), Pearson correlation among replicates (minimum, maximum), sequencing instrument, library type (SR=single read or PE=paired-end read, read length (nucleotides), sequencing type = polyA/total).

The degree of homogeneity among samples, as measured by Pearson’s correlation coefficient, was lowest for the Zhang dataset followed by GTEx (the estimated biological coefficients of variation shown in Figure A1-panel A of Additional File 1 also support this fact). As expected, the other datasets had replicates that are more homogeneous because they were obtained from inbred animals or cultured cell lines, in contrast to the GTEx or Zhang datasets containing tissues for different human individuals. For the Zhang and NGP Nutlin datasets, lncRNAs showed relatively higher heterogeneity across samples than mRNAs. In addition, lncRNAs showed on average lower expression than mRNAs (Figure A1-panel B).

### Comparison of normalization methods

Comparing DE tools requires careful attention to the normalization methods. Previous studies^17,20,42,43^ have pointed out that the normalization procedure can affect the result of DE analysis. We therefore first explored the performance of 5 normalization methods that are used in conjunction with the DE methods evaluated in this study. The normalization methods were compared using the metrics from Dillies et al.^42^, such as their capability to reduce variability that attributes to technical sources, their capability of eliminating bias due to library size differences, and their effect on DGE analysis.

Box plots of log transformed counts showed that for all 6 datasets all normalization methods succeeded in aligning the sample specific distributions and hence no library size effects were noticeable after normalization (Figures A3-A8 of Additional File 1). Furthermore, the condition-specific genewise coefficient of variation (CV), which is a proxy for intra-group biological variability, was reduced for all datasets by all normalization procedures (Figures A9-A14). Nearly equal levels of biological variability across methods were observed, even with quantile normalization that was found to result in high CV in other studies^42,44^. The overlap of DE genes with different normalization methods was high (Figure 2). Ignoring quantile normalization (QN), a minimum of 96% similarity was observed. QN-based DE analysis gives deviating results, particularly for designs with small numbers of replicates (< 5); the lowest proportion of similarity was 68%. Overall, the results suggest that all normalization methods perform almost equally, except QN. Nevertheless, for the concordance analysis of the DE tools (see next section) we include a limma method that uses QN (named limmaQN) to further investigate its effect on other relevant performance metrics of DE tools.

**Figure 2.**
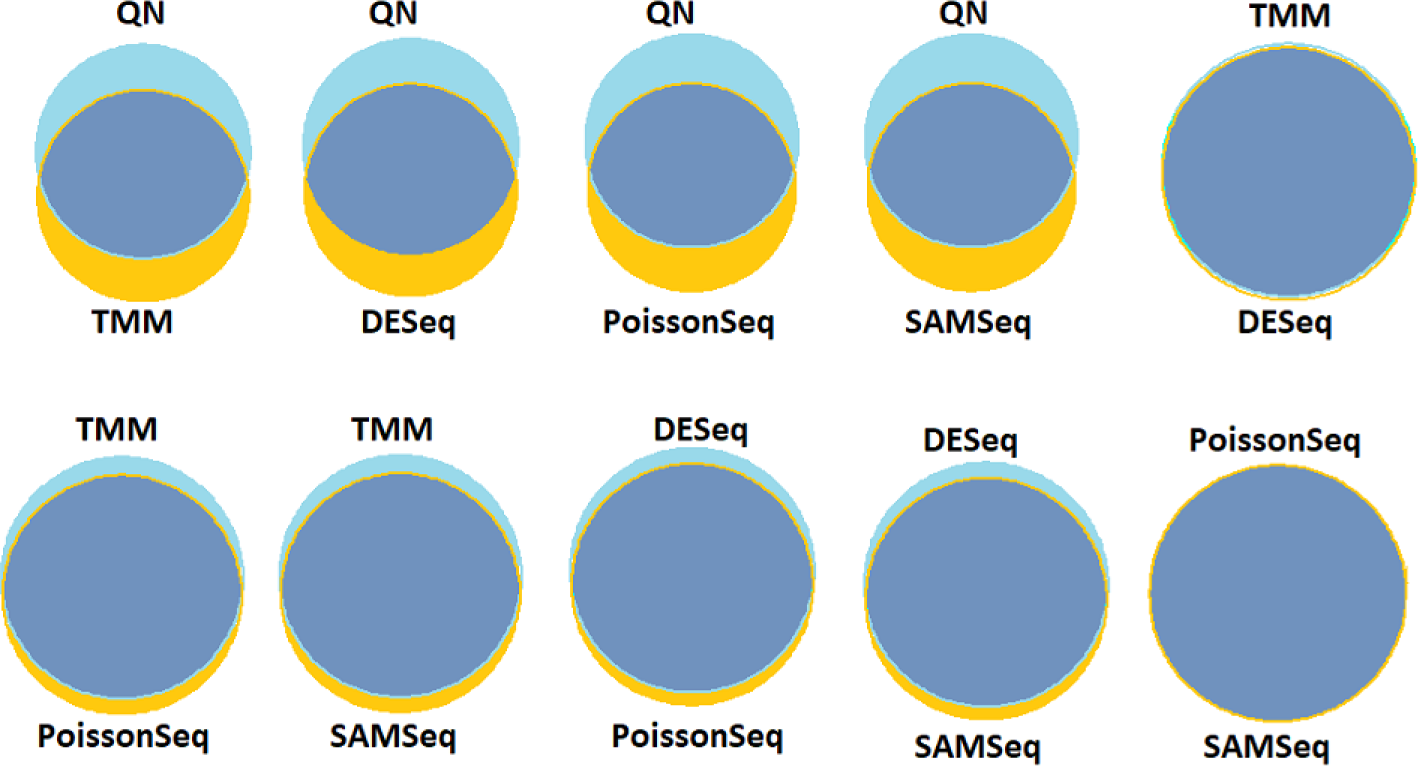
Effect of normalization methods on DGE analysis. The Venn diagrams show the level of concordance between pairs of differential gene expression (DGE) analyses each using different normalization methods but the same statistical test (moderated t-test from *limma* package) for the CRC AZA data. The size of the circles is proportional to the number of significant DE genes (at 5% FDR) using the particular normalization. All considered normalization methods generally show strong concordance except for quantile normalization. TMM and DESeq normalization as well as PoissonSeq and SAMSeq normalization showed the strongest similarity. The values underlying these Venn diagrams are presented in Additional Files 1 Table A2.

### Concordance Analysis

Fourteen DE tools were run on 6 RNA-seq datasets, and (dis)similarities among the results were examined. The concordance analysis focused on four quantitative and one qualitative metric: (1) number of genes identified as significantly differentially expressed (SDE); (2) the degree of agreement on gene ranking; (3) similarity of fold-change estimates; (4) handling of genes with special characteristics (lncRNAs, genes with low counts, genes with outliers), and (5) computation time. The results for individual datasets are presented in Additional File 1 Sections 3.3-3.8.

Results show that the 14 DE tools show substantial variability in the numbers of SDE genes. The marginal summary across all datasets indicates that DESeq, baySeq, and limmaQN detected the smallest number of SDE genes, whereas QuasiSeq and SAMSeq returned the largest numbers (Figure 3, Figure A15). The variability among DE tools with respect to the number of SDE genes seems to be related to the biological variability in the dataset. For the Zhang and GTEx RNA-seq datasets, characterized by the largest intra-group biological variability, the numbers of SDE genes were quite different among the DE tools. In contrast, the numbers of SDE genes from the NGP Nutlin and CRC AZA datasets, all displaying low biological variability, were relatively similar among tools. lncRNAs and low abundant genes in general were underrepresented among the SDE genes (Additional File 1 Section 3.3-3.8). For example, 25% of the SDE genes were lncRNAs, whereas the data contains 40% lncRNAs.

**Figure 3.**
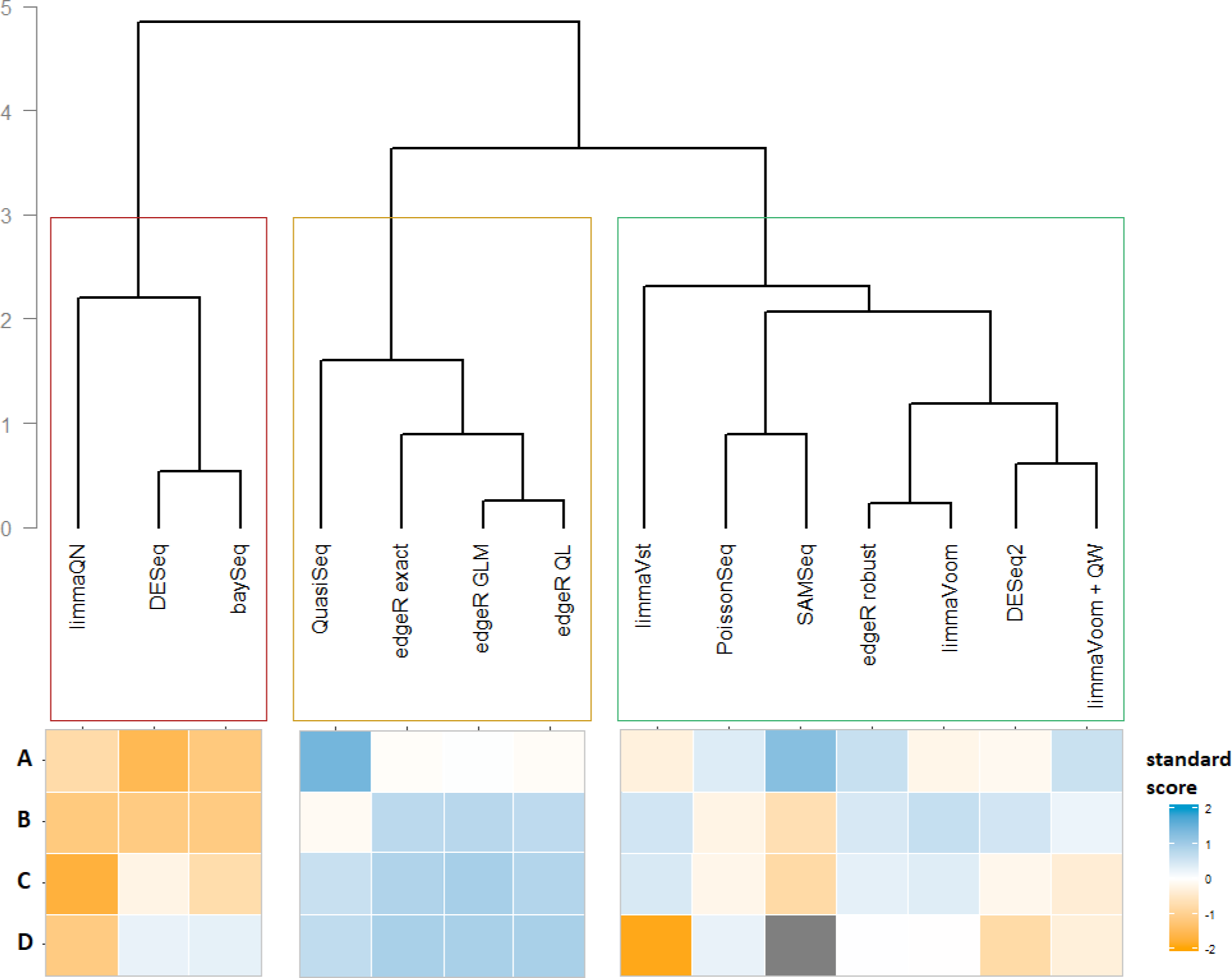
Summary of concordance analysis results. Hierarchical clustering of 14 DE tools based on standard scores of comparison metrics (A. number of significantly differentially expressed (SDE) genes detected at 5% FDR, B. overlap among tools in detecting SDE genes at 5% FDR, C. gene ranking agreement, and D. similarity of log fold change (LFC) estimates) marginalized across the 6 datasets. First, observed values of concordance metrics (proportions and correlations) for each DE tool from a given dataset are converted to standard scores, (value - mean)/standard deviation, and then the average of standard scores of each DE tool across datasets are presented. A negative value, for example, for number of SDE genes indicates that the number of SDE genes detected by the tool is lower than the average across all the 14 tools. Afterwards, the euclidean distance among the marginal standardized scores of DE tools for the four comparison metrics are computed and complete linkage method of agglomerative clustering is applied. The methods are grouped in three clusters. The heatmap plot underneath the cluster shows the individual marginal scores of each DE tool for the four concordance measures. Since fold-change estimate from SAMSeq is in terms of sum of ranks, it was excluded from comparison of LFC estimates, and hence it is indicated by gray color in the heatmap plot.

Many DE tools detected the same set of SDE genes (Figure 3, Figure A15). On average, limmaQN, DESeq, baySeq, and SAMSeq showed the smallest concordance with all other tested DE tools. It was also observed that the overlap of SDE is smaller for lncRNAs than for mRNAs (Figure A16). In the Zhang dataset, there is less than 70% and 60% SDE overlap across all DE tools for mRNAs and lncRNAs, respectively.

Accurate gene ranking is an essential step for downstream analysis such as gene set enrichment analysis (GSEA)^45^. In the current study, genes are ranked taking into account both the significance and magnitude of differential expression, using the n-score^46^. Summarized results across datasets (Figure 3, Figure A15) indicate that all DE tools strongly agree, except for baySeq, SAMSeq, and limmaQN. Apart from baySeq, this is somewhat in contrast to the findings in Soneson et al.^18^. This might be due to the difference in the score used to rank genes, as only P-values were used to rank genes in Soneson et al.^18^. Except for limmaQN, gene ranking agreement among all DE tools was nearly the same for lncRNAs and mRNAs (Figure A16). A slightly lower agreement for lncRNAs was observed when the most variable dataset (Zhang) is used. limmaQN consistently exhibited the lowest level of agreement with all other DE tools with respect to the concordance metrics discussed so far, and this is likely due to the quantile normalization it uses (see higher).

For moderately to highly expressed mRNA genes (the 75% highest expressed genes) the log-fold-change (LFC) estimates from all DE tools were strongly correlated, with Pearson correlation coefficients ranging from 0.95 to 0.99 (Additional File 1, Sections 3.3-3.8). For the 25% lowest abundant genes, the estimates show weaker correlations (Pearson correlation less than 0.8 in most cases). Similarly, the correlations for lncRNAs were lower than for mRNAs (Figure A16).

In Section 3.2.1 of Additional File 1, we also examined the handling of genes with unusual counts across samples. It turned out that only the limma tools, DESeq2, and baySeq showed concordant decisions with edgeR robust and SAMSeq (particularly developed to better cope with outliers), suggesting that they are neither sensitive nor conservatives tools.

The computation time to run DGE analysis presented in Section 3.2.2 of Additional File 1 shows that baySeq and DESeq require the longest time, whereas limma tools and PoissonSeq run fast. For RNA-seq data with 10 replicates per group and 19,150 mRNAs, the slowest tools, baySeq and DESeq, were approximately 8,000 and 2,000 times slower than the fastest tool, limmaQN, respectively.

In an attempt to come to an overall conclusion, the results were combined in a hierarchical clustering analysis of the DE tools resulting in 3 clusters (Figure 3). DESeq, baySeq, and limmaQN cluster together, generally showing the least number of SDE genes, lower overlap, and lower gene ranking agreement with all other tested DE tools. The second cluster includes QuasiSeq, edgeR exact, edgeR GLM, and edgeR QL, showing the highest concordance with respect to calling SDE, gene ranking, and LFC estimates. This cluster also shares the same modeling scheme (negative binomial models) and normalization method (TMM). The third cluster includes DESeq2, edgeR robust, limmaVoom(+QW), limmaVst, PoissonSeq, and SAMSeq, showing high concordance to all other DE tools like the second group except that they showed slightly less similar LFC estimates. Moderating the LFC estimate for low expressed and/or outliers genes (DESeq2 and edgeR robust) or transformation of counts to log scale (limmaVoom (+QW) and limmaVst) may explain slightly different LFC estimates.

### Simulation results

The non-parametric *SimSeq*^47^ procedure was applied to realistically simulate RNA-seq expression data. The simulation technique involves sub-sampling of replicates from a real RNA-seq dataset with a sufficiently large number of replicates. In this way, the underlying characteristics of the source dataset are preserved, including the count distributions and variability. Three series of simulations were performed, each starting from a different RNA-seq source dataset: Zhang, NGP Nutlin, and GTEx data. The degree of homogeneity among the replicates in these datasets varies, reflecting different levels of intra-group biological variability (Table 2 and Figure A1). The Zhang and NGP Nutlin datasets include annotated lncRNAs along with mRNAs, whereas the GTEx RNA-seq dataset contains only annotated mRNA genes. Therefore, simulated counts for mRNA and lncRNA are sampled from mRNA and lncRNA counts of the source dataset, respectively.

Gene expressions were simulated under a wide range of scenarios that may affect the performance of DE tools: different numbers of replicates ranging from 2 to 40, different proportions of true DE genes (0 to 30%), two gene biotypes (mRNA and lncRNA), and different levels of intra-group biological variability (as present in the three source datasets). From the simulation results, the actual false discovery rate (FDR), true positive rate (TPR), and false positive rate (FPR) were computed for 13 DE tools. The comparison between the two gene biotypes was done in two ways: simulating lncRNA data only or simulating lncRNA and mRNA jointly, but analyzing separately.

#### False discovery rate

FDR refers to the average proportion of incorrect discoveries among SDE genes. It refers to DE methods applied at a user-defined nominal level of the FDR, which is here set to 5%. A good DE tool has actual FDR close to the nominal level, and has high TPR. Our results for simulations starting from the Zhang dataset (Figure 4) indicate that the FDR is not controlled by all DE tools. Only PoissonSeq, limmaVoom, limmaQN, limmaVst, SAMSeq, and DESeq gave acceptable actual FDR levels. The other methods showed severe FDR inflation, particularly when only a small fraction of genes is DE (Figure 4 and Figures A54 and A55 of Additional File 1). FDR levels may even exceed 50%, which means that more than half of the SDE called genes may be false discoveries. For most DE tools, the FDR control was better with increasing number of replicates. The gene biotype was also influential, with considerably poorer performance for lncRNAs. When starting from the NGP Nutlin source dataset, the results were better (Figure 5 and Figure A63), with good FDR control for all DE tools, even with small sample sizes. Only for simulations with 5% of true DE genes, the FDR control was lost (Figure A63). The difference in FDR control between the Zhang and NGP Nutlin datasets is likely explained by their intra-group variability (see Table 2, Figure A1). In line with this, GTEx has intermediate homogeneity and the FDR control in simulations is somewhere in between the extreme Zhang and NGP Nutlin datasets (Figure A65). In Section 4.5.1 of Additional File 1, we separately demonstrated the actual FDR and TPR of DE tools for low and high abundant mRNAs based on a simulation starting from the GTEx data. In general, the tools showed relatively worse performance for low abundant mRNAs (high FDR and low TPR) as seen for lncRNAs.

**Figure 4.**
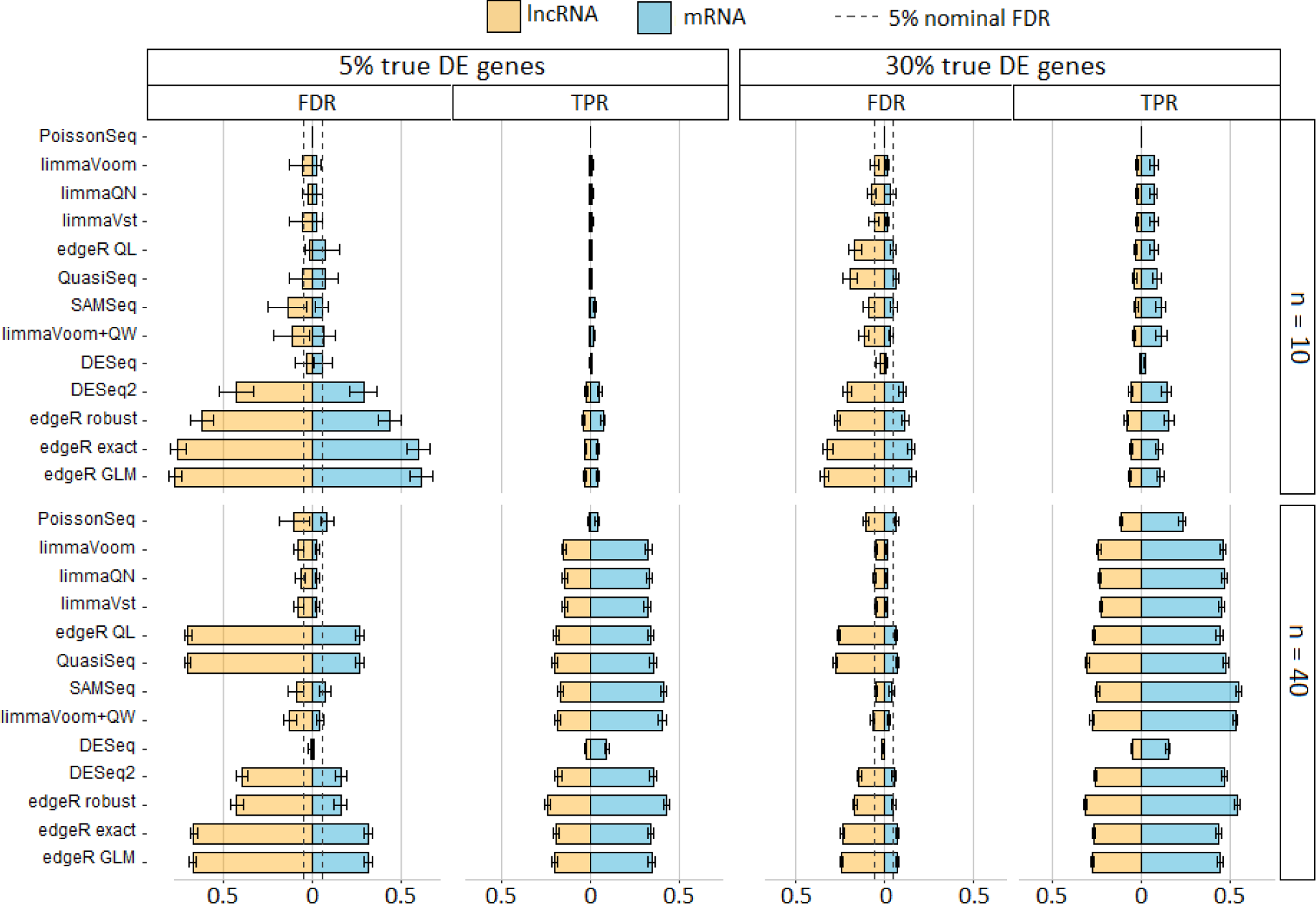
False discovery rate and true positive rate of DE tools using simulated data from the Zhang RNA-seq dataset. It shows the actual FDR and TPR (at 5% nominal FDR) of 13 DE tools from joint simulation and DGE analysis of mRNA and lncRNA. These particular results are from simulations with 5% and 30% true DE genes among 10,000 genes (constituting approximately 30% lncRNAs and 70% mRNAs) for designs with n = 10 and 40 replicates per group. The error bars indicate the 95% confidence interval estimate. Gray dashed line is included for FDR results to indicate the nominal FDR level (5%). Number of replicates and proportion of true DE genes have higher impact on the TPR and FDR of DE tools. Although negative binomial models (edgeR, DESeq2, and QuasiSeq) showed higher sensitivity, in general they showed higher and uncontrolled FDR for simulated datasets with lower number of replicates and lower proportion of true DE genes. In contrast, among discrete distribution models, DESeq and PoissonSeq showed better capability of controlling FDR, with actual FDR below the threshold level (5%), but these tools have lower sensitivity than all other DE tools. SAMSeq and limma tools consistently showed fair level of true FDR with comparable TPR to negative binomial models. In most cases, DE tools exhibited higher FDR and lower sensitivity for lncRNAs than for mRNAs.

**Figure 5.**
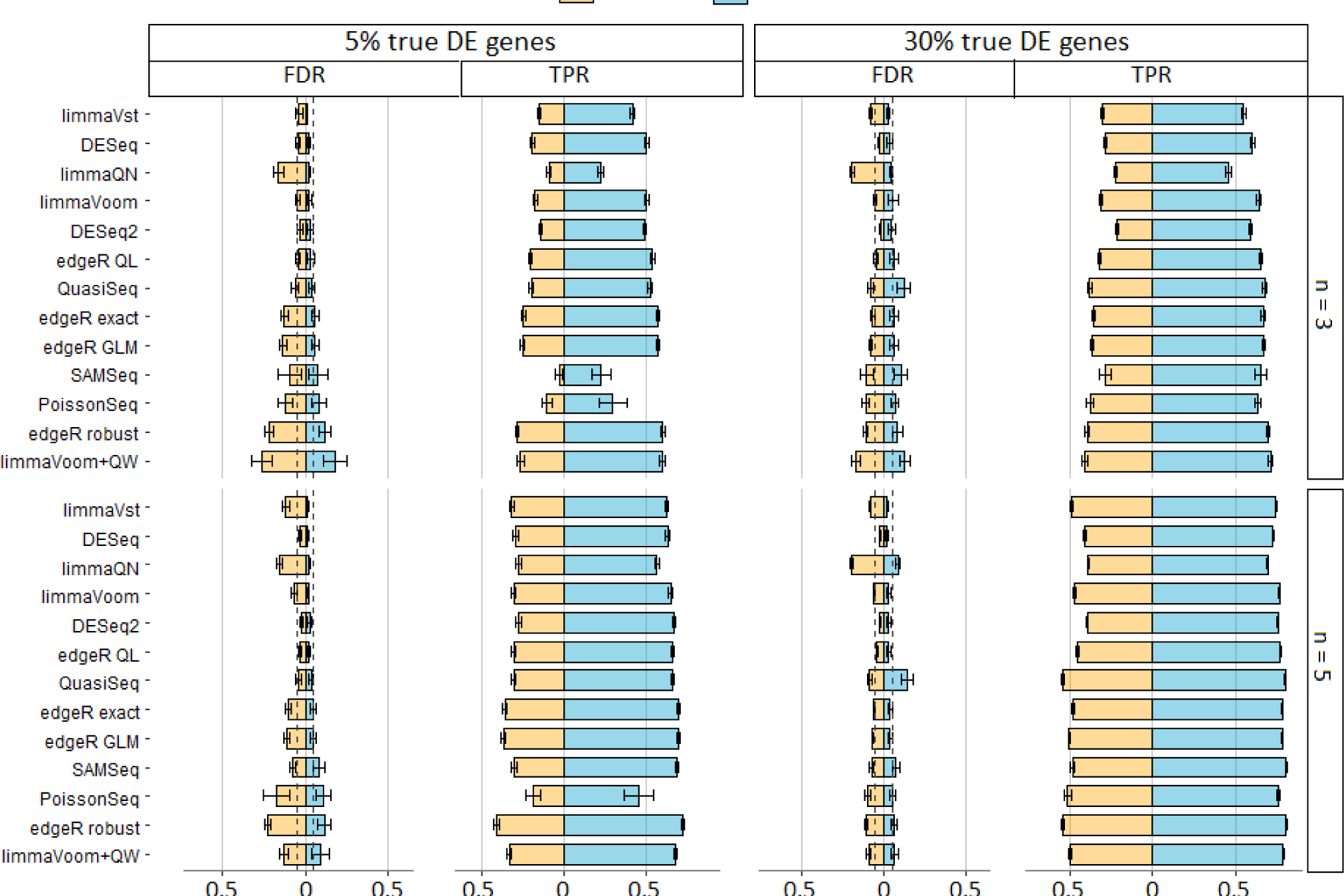
False discovery rate and true positive rate of DE tools using simulated data from the NGP Nutlin datasets. It shows the actual FDR and TPR (at 5% nominal FDR) of 13 DE tools from joint simulation and DGE analysis of mRNA and lncRNA expression data. These particular results are from simulations with 5% and 30% true DE genes among 10,000 genes (constituting approximately 35% lncRNAs and 65% mRNAs) for designs with n = 3 and 5 replicates per group. The error bars indicate the 95% confidence interval estimate. Gray dashed line is included for FDR results to indicate the nominal FDR level (5%). The performance of DE tools for gene expression data simulated from the NGP Nutlin dataset is better due to the lower intra-group biological variability in the simulated dataset, which is reflected from the source data. All DE tools showed acceptable level of actual FDR with sensitivity much better for designs with 3 to 5 number of replicates and 30% true DE genes. DE tools’ TPR again appeared to be lower for lncRNAs than for mRNAs. DESeq and PoissonSeq also showed improved sensitivity for this simulation.

#### True positive rate

The TPR, which is also known as sensitivity, is the average proportion of SDE genes among the true DE genes. The TPR should be sufficiently large, otherwise one cannot expect to find many of the true DE genes. Our simulation results indicated that the TPR increases with increasing number of replicates (Figure 4, Figure A58, and Figure A63). PoissonSeq and DESeq showed substantial smaller TPR than the other tools, even in simulations with more than 20 replicates per group. Because of the trade-off between FDR and TPR, a large TPR is expected for DE tools with a large actual FDR. This was observed for edgeR tools, QuasiSeq, and DESeq2. The tools that showed better capability of controlling FDR (limma tools and SAMSeq), also had TPR nearly equal to the TPR from liberal tools. The results also showed that a slightly higher TPR was obtained when more DE genes are present (Figures A56 and A57), and that the sensitivity for detecting DE mRNA was larger than for lncRNA. Similar conclusions can be drawn from the simulations using the more homogeneous datasets; NGP Nutlin and GTEx simulations result in higher TPR (Figure 5 and Figures A63 and A65 in Additional File 1).

Finally, analyzing simulated counts using the default filtration setting of DE tools (particularly DESeq2 and PoissonSeq) resulted in slightly improved performance at a cost of losing a considerable proportion of lncRNAs (Figure A64 of Additional File 1).

#### False positive rate

The FPR refers to the probability of calling a gene SDE in a scenario with no DE genes at all. Results are shown in Figure A53 of Additional File 1. They demonstrate that all DE tools resulted in a FPR of less than 1%. The results were similar for both gene biotypes (mRNAs and lncRNAs), except for a slightly higher FPR for lncRNAs than for mRNAs. The FPR was generally larger for methods relying on the negative binomial distribution. This finding is in line with conclusions from a previous comparative study^17^, in which it was concluded that the number of false predictions of differential expression from DE tools (majority of these DE tools are also the part of our study) is sufficiently low even for genes with low counts (gene expressed in the lower 25%).

#### Simulation of lncRNA expression data only

Results presented up to this point came from simulating, normalizing and analyzing lncRNAs and mRNAs together. Of note, joint analysis of the two gene biotypes may affect results. For example, estimates of gene specific dispersion parameters for negative binomial models are often done by sharing information across all genes using empirical Bays strategy^26,32,37^, and hence the results for lncRNAs depend on mRNA read counts and vice versa. In addition, adjusted P-values aimed at controlling FDR are calculated taking into account the total number of genes included in the analysis^48^. Therefore, we also evaluated the performance of the DE tools with only lncRNA data, using the same simulation procedures. Our conclusions remain the same. The results are shown in Figure A62 of Additional File 1. The FDR control is generally worse when analyzing lncRNA separately, particularly for small replicate sizes. Only a small reduction in TPR is observed.

### Web application

All simulation results can be consulted and visualized with a web application: http://statapps.ugent.be/tools/AppDGE/.

## Conclusions

Our study evaluated the performance of widely used statistical tools for testing differential gene expression (DGE) in RNA-seq data, with separate analysis of mRNA and lncRNA. Several gene expression studies indicated that the expression of the majority of lncRNAs is characterized by low abundance^2,10,12,13^, high noise^11^, and tissue-specific expression^10^. These characteristics are very challenging for DE tools, and may potentially negatively affect tool performance^14,15^.

Concordance analysis of the DE tools across 6 diverse RNA-seq datasets revealed that the DE tools lack strong consensus on identifying a set of significantly differentially expressed (SDE) genes, gene ranking, fold-change estimates, and handling low count genes. For datasets with a small number of replicates and/or heterogeneous replicates, the disagreement is even worse. Lower concordance was observed for lncRNAs than for mRNAs. In particular, limmaQN, baySeq, and DESeq (also PoissonSeq and SAMSeq to a smaller extent) showed in general lower concordance with other DE tools and also exhibited data dependent performance. In contrast, edgeR, DESeq2, limmaVoom(+QW), and limmaVst showed better agreement with other DE tools and consistent characteristics across datasets.

Results of our large non-parametric simulation study revealed that there are substantial differences among methods with respect to FDR control and sensitivity. The DE tool performance is strongly affected by the sample size, biological variability, and biotype of the transcripts. FDR control at the nominal level is good for all methods for datasets with small biological variability, even with only 5 biological replicates per condition. On the other hand, for datasets that are more variable, the FDR control is only guaranteed for larger sample sizes, and only with the following methods: PoissonSeq, limma tools, SAMSeq and DESeq. All other DE methods result in actual FDR levels far above the nominal level, up to an FDR exceeding 50% for lncRNAs even when studying 40 replicates per condition. Differences among tested tools in terms of sensitivity are not very large, except for PoissonSeq and DESeq, and limmaQN to a smaller extent, which showed the lowest sensitivities among all methods. For highly variable data, a maximum sensitivity of 50% for mRNAs is obtained with 40 replicates per condition. For homogeneous samples, this level of sensitivity is already reached with 4 and 5 replicates per condition.

In the light of promoting reproducible science, it is essential to select a DE tool that succeeds in controlling the FDR level under a large range of conditions. Among these DE tools, one can select one with a high sensitivity. If a DE tool has an actual FDR far larger than the nominal level, many of the discoveries claimed will be false discoveries. If one is prepared to allow for a large proportion of false discoveries, it is generally better to still use a DE tool with good FDR control, but to apply the method at a larger nominal FDR level. In this way, the researcher still controls the error rate. This reasoning implies that the selection of a DE tool may never rely purely on the sensitivity. High sensitivities may be expected from methods with a large actual FDR (and hence poor FDR control). These high sensitivities are illusive in the light of the large proportion of false discoveries.

Combining all results, we conclude that limma with variance stabilizing transformation, limma voom (with and without quality weighting), and SAMSeq control the FDR reasonably well, while not sacrificing sensitivity. These methods end up in the same cluster based on the concordance analysis.

## Methods

### RNA-seq datasets

A total of 6 datasets are used in this study. All of them are publicly accessible. The Zhang dataset, containing 498 neuroblastoma tumors, was retrieved from Zhang et al.^41^ (GEO accession number GSE49711). In short, unstranded polyA+ RNA sequencing was performed on the HiSeq 2000 instrument (Illumina). Paired-end reads with a length of 100 nucleotides were obtained. To quantify the full transcriptome, raw fastq files were processed with Kallisto^49^ v0.42.4 (index build with GRCh38-Ensembl v85). The pseudo-allignment tool Kallisto was chosen above other quantification methods as it is performing equally good but faster. For this study, a subset of 172 patients with high-risk disease were selected, forming two groups: the MYCN amplified (n=91) and MYCN non-amplified (n=81) tumors. Randomly selected 20 samples from each group were used to study concordance analysis of the DE tools, whereas all samples were considered for the simulation study.

Second, the NGP Nutlin-3 dataset (GEO accession number GSE104756) contains 10 biological replicates of NGP neuroblastoma cells grown in 6-well culture flasks treated with either nutlin-3 (8μM racemic mixture) or vehicle (ethanol). RNA was extracted using the RNeasy mini kit (Qiagen), quantified using Nanodrop-100, and quality controlled using FragmentAnalyzer (High Sensitivity RNA Analysis Kit, DNF-472-0500, Advanced Analytical). 100 ng of total RNA was used for TruSeq stranded mRNA library prep (Illumina), followed by paired-end sequencing (2x75 nucleotides) on a NextSeq 500 instrument (Illumina). On average, 20 million read pairs per sample were generated. Raw fastq files were processed with Kallisto2 v0.42.4 (index build with GRCh38-Ensembl v85).

From the Gene Expression Omnibus (GEO) repository of analysis-ready RNA-seq dataset^50^, we accessed two datasets: Hammer (GEO accession number GSE20895) and Bottomly mRNA-seq datasets (GEO accession number GSE26024). The Hammer mRNA-seq dataset contains the results of a study on the L4 dorsal root ganglion (DRG) of rats with chronic neuropathic pain induced by spinal nerve ligation (SNL) of the neighboring (L5) spinal nerve (two controls and two with L5-SNL induced chronic neuropathic pain)^38^. The study includes RNA-seq samples of each treatment at two different time points, and in our study, we only used samples from the first time point. The Bottomly mRNA-seq data is an expression set obtained from the brain striatum tissues of two mice strains (it included 10 biological replicates of the C57BL/6J (B6) strain and 11 biological replicates of the DBA/2J (D2) strain)^39^. Details about library preparation and features of these datasets can be obtained from the original papers cited above.

We also used a subset of RNA-seq dataset from the GTEx (Genotype-Tissue Expression) project^40^, which provides a large database of human gene expression data with more than 3000 RNA-seq samples from 54 different tissues. For our study, we selected all biological replicates of hippocampus (n1=28) and hypothalamus (n2=30) mRNA-seq counts. All information about this dataset can be obtained from the GTEx project database. Randomly selected 20 samples from each tissue were used for concordance analysis of the DE tools and all samples were considered for the third simulation study.

The colorectal cancer (CRC AZA) RNA-seq dataset contains 3 biological replicates of HCT-116 cells grown in 6-well culture flasks treated with either Azacitidine (1μM) or vehicle (DMSO). RNA was extracted using the RNeasy mini kit (Qiagen), quantified using Picogreen, and quality controlled using Bioanalyser FragmentAnalyzer. 500 ng of total RNA was used for TruSeq stranded mRNA library prep (Illumina), followed by paired-end sequencing (2x100 nucleotides) on a HiSeq instrument (Illumina). On average, 40 million read pairs per sample were generated. Raw fastq files were processed with Bowtie2/Cufflinks (index build with GRCh37-Ensembl v75). The count data can be accessed in Additional File 2.

### DE tools and normalization methods

Five normalization methods are evaluated in this study: quantile normalization (QN)^31^ implemented in limma (limmaQN), Trimmed Mean of M-values (TMM)^25^ implemented in edgeR, limma (limmaVoom and limmaVoom+QW), baySeq, and QuasiSeq, Medians of Ratios (MR)^30^ implemented in DESeq, DESeq2, and limma (limmaVst), the goodness-of-fit statistics^36^ approach in PoissonSeq, and the re-sampling technique^23^ implemented in SAMSeq. To examine the effect of normalization on DGE analysis, we applied moderated t-test of the *limma* package in combination to each of the five normalization methods. Afterwards, the extent of dissimilarity of results is used as an indicator of normalization effect on DGE analysis.

The fourteen DE tools that are considered in this paper are summarized in Table 1. All methods are available as R software packages^16^. Most methods were applied using their default settings, except for DESeq2 and PoissonSeq, which required certain settings to be changed to avoid filtering out too many low abundant genes (such as lncRNA). In particular, in DESeq2 the independent filtering was disabled in the results function, and in PoissonSeq the filter cutoffs for the sum and average counts of genes across samples were set to 1 and 0.01, respectively. All DE tools were applied at the nominal 5% FDR level, unless mentioned otherwise. All computations were performed in R version 3.3.2^16^. baySeq was not included in the simulation study due to its slow computation time. The reader can also find the R scripts in the Additional Files 2.

### Concordance analysis

The number of SDE genes detected for each dataset are compared among 14 DE tools. For the Zhang and NGP Nutlin datasets, we explored the proportion of mRNAs and lncRNAs within the set of SDE genes. For the other datasets, we explored the proportion of low and high expressed mRNAs among the set of SDE mRNAs. In particular, we divided the genes into 4 equally large groups based on the quartiles of their average normalized read counts across samples (DESeq normalization). To study the concordance in SDE calling between any two DE tools, we calculated the extent of overlap as the proportion of SDE genes commonly detected by the two tools. To rank genes in the order of significance of differential expression, taking into account the biological importance of the significance we calculated the n-score^46^,

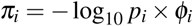

where *ϕ_i_* is the absolute value of the estimated LFC for the *i^th^* gene and *p_i_* is the raw P-value, except for SAMSeq, for which *p_i_* is replaced by 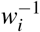, where *w_i_* is the Wilcoxon statistic^23^. For baySeq, *p_i_* is the estimated posterior probability of differential expression (estimated Bayesian False Discovery Rate, BFDR)^?^. Afterwards, we used Spearman’s rank correlation to evaluate the degree of agreement among DE tools with respect to gene ranking.

### Non-parametric simulation methods

SimSeq^47^ simulates a matrix of RNA-seq read counts by sub-sampling samples from a source RNA-seq dataset. It creates a set of DE genes based on prior information from analyzing the source dataset (which has larger number of replicates than the simulated expression data). Details about the method are given in Benidt et al.^47^ and in Section 4 of Additional File 1. Results from DGE analysis of the source RNA-seq dataset is available in Table A4 of Additional File 1. Three sets of simulations were performed, each starting from a different source dataset: the Zhang, NGP Nutlin, and GTEx datasets. Gene expressions are simulated containing a given proportion of true DE genes (ranging from 0 to 30%) from a total of 10,000 genes. Each gene in every simulated count matrix has a total of at least one expression in each group. Also, counts are simulated for varying numbers of replicates per group (2 to 40, 2 to 5, and 2 to 14 for simulations that start from the Zhang, NGP Nutlin, and GTEx datasets, respectively). For joint simulation of mRNAs and lncRNAs (for Zhang and NGP Nutilin), each simulated count matrix includes a number of lncRNAs and mRNAs in the set of true DE genes and non-DE genes, proportional to their numbers in the source dataset (for the Zhang data 70% mRNA and for NGP Nutlin data 65% mRNA). We ran 100 independent simulations for each scenario. On each simulated dataset, we applied the various DE tools and calculated P-values, adjusted P-values, LFC, and other relevant statistics. These statistics were subsequently used for computing the comparative performance metrics.

For a given fraction of true DE genes, the SimSeq method generates data for which it is known which genes are truly DE. To each simulated dataset all DE tools are applied and genes are called SDE if their FDR adjusted P-values are smaller than the nominal FDR level (5%, unless mentioned otherwise). For a single simulated dataset the observed false discovery proportion *(FDP)* is calculated as *FDP = FP/(FP + TP)*, where *FP* and *TP* are the numbers of incorrect (false) and correct (true) rejection of the null hypothesis, respectively. The actual False Discovery Rate (FDR) is then computed as the average of the *FDPs* over all simulations. When the FDR is computed for a scenario with no DE genes at all, the FDR is generally known as the false positive rate (FPR). For this scenario, the SimSeq procedure simply samples replicates from only one group of the source dataset and randomly assigns the samples to one of two arbitrary mock groups. The actual True Positive Rate (TPR), which is also called sensitivity, is defined as the average of the true positive proportions (*TPP*) across independent simulations, *TPP = TP/(TP + FN)*, where *FN* is the number of incorrect decisions to fail rejecting the null hypothesis.

The quality of simulated data was examined using the quality metrics proposed by Soneson and Robinson^51^ and implemented by their countsimQC R package (version 0.5.2). The metrics evaluate the average expressions of genes, variability, mean-variance relationship, correlations among replicates, correlations among genes, and fractions of zero counts. The quality assessments were positive in all aspects, and the reports generated by the countsimQC R package can be found in Additional File 2.

### Web application

A web application is developed, using the R Shiny package (version 1.0.5)^52^. This user interface allows the user to consult and visualize all detailed results from this paper. It can be reached at http://statapps.ugent.be/tools/AppDGE/.

## Additional Files

Additional results (tables, figures, and discussion) are compiled in a separate file with name **Additional File 1. Additional File 2** includes R source codes used in this study, reports for quality assessment of simulated data, and CRC AZA count dataset. It is accessible through a github repository with name “Additional-File-2” (https://github.com/CenterForStatistics-UGent/Additional-File-2.git). We created a web tool that assists researches to choose the best DE tools for a given RNA-seq study based on our simulation based comparison of 13 DE tools. It can be accessed with a link http://statapps.ugent.be/tools/AppDGE/.

## Author’s contributions

O.T., J.V., and P.M. conceived and supervised the study, A.A. analyzed and reported the results, K.P. prepared the initial report, A. A. made the final draft and the web took, C.E. prepared the data. All authors reviewed and approved the final version of the manuscript.

## Acknowledgments

We are grateful to Janssen Pharmaceutica (Belgium) for providing the cell line CRC AZA RNA-seq data. We are also grateful to Matthias Fischer for providing the raw Zhang data, Steve Lefever for creating Figure 1 (the general workflow of this study).

## References

1. Derrien, T. et al. The GENCODE v7 catalog of human long non-coding RNAs: analysis of their gene structure, evolution, and expression. Genome research 22, 1775–1789 (2012).

2. Volders, P.-J. et al. LNCipedia: a database for annotated human lncRNA transcript sequences and structures. Nucleic acids research 41, D246–D251 (2013).

3. Maass, P. G., Luft, F. C. & Bahring, S. Long non-coding RNA in health and disease. J. Mol. Medicine 92, 337–346 (2014). URL http://dx.doi.org/10.1007/s00109-014-1131-8. DOI 10.1007/s00109-014-1131-8.

4. Gong, J., Liu, W., Zhang, J., Miao, X. & Guo, A.-Y. lncRNASNP: a database of SNPs in lncRNAs and their potential functions in human and mouse. Nucleic acids research 43, D181–D186 (2015).

5. Trypsteen, W. et al. Differential expression of lncRNAs during the HIV replication cycle: an underestimated layer in the HIV-host interplay. Sci. Reports 6 (2016).

6. Wallaert, A. et al. Long noncoding RNA signatures define oncogenic subtypes in T-cell acute lymphoblastic leukemia. LEUKEMIA (2016). URL http://dx.doi.org/10.1038/leu.2016.82.

7. Gutschner, T. & Diederichs, S. The hallmarks of cancer: a long non-coding RNA point of view. RNA biology 9, 703–719 (2012).

8. Wang, Z., Gerstein, M. & Snyder, M. RNA-seq: a revolutionary tool for transcriptomics. Nat. Rev. Genet. 10, 57–63 (2009).

9. Cabili, M. N. et al. Integrative annotation of human large intergenic noncoding RNAs reveals global properties and specific subclasses. Genes & development 25, 1915–1927 (2011).

10. Tsoi, L. C. et al. Analysis of long non-coding RNAs highlights tissue-specific expression patterns and epigenetic profiles in normal and psoriatic skin. Genome biology 16, 1 (2015).

11. Kornienko, A. E. et al. Long non-coding RNAs display higher natural expression variation than protein-coding genes in healthy humans. Genome biology 17, 14 (2016).

12. Ren, H. et al. Genome-wide analysis of long non-coding RNAs at early stage of skin pigmentation in goats (Capra hircus). BMC genomics 17, 1 (2016).

13. Xia, J. et al. Characterization of long non-coding RNA transcriptome in high-energy diet induced nonalcoholic steatohepatitis minipigs. Sci. Reports 6 (2016).

14. Love, M. I., Huber, W. & Anders, S. Moderated estimation of fold change and dispersion for RNA-seq data with DESeq2. Genome Biol. 15, 550 (2014). DOI 10.1186/s13059-014-0550-8.

15. Raithel, S. et al. Inferential considerations for low-count RNA-seq transcripts: a case study on the dominant prairie grass andropogon gerardii. BMC genomics 17, 140 (2016).

16. R Development Core Team. R: A Language and Environment for Statistical Computing. R Foundation for Statistical Computing, Vienna, Austria (2008). URL http://www.R-project.org. ISBN 3-900051-07-0.

17. Rapaport, F. et al. Comprehensive evaluation of differential gene expression analysis methods for RNA-seq data. Genome biology 14, 1 (2013).

18. Soneson, C. & Delorenzi, M. A comparison of methods for differential expression analysis of RNA-seq data. BMC bioinformatics 14, 1 (2013).

19. Schurch, N. J. et al. How many biological replicates are needed in an RNA-seq experiment and which differential expression tool should you use? RNA (2016).

20. Bullard, J. H., Purdom, E., Hansen, K. D. & Dudoit, S. Evaluation of statistical methods for normalization and differential expression in mRNA-seq experiments. BMC bioinformatics 11, 94 (2010).

21. Seyednasrollah, F., Laiho, A. & Elo, L. L. Comparison of software packages for detecting differential expression in RNA-seq studies. Briefings bioinformatics 16, 59–70 (2015).

22. Sahraeian, S. M. E. et al. Gaining comprehensive biological insight into the transcriptome by performing a broad-spectrum RNA-seq analysis. Nat. Commun. 8 (2017).

23. Li, J. & Tibshirani, R. Finding consistent patterns: a nonparametric approach for identifying differential expression in RNA-seq data. Stat. methods medical research 22, 519–536 (2013).

24. Robinson, M. D. & Smyth, G. K. Small-sample estimation of negative binomial dispersion, with applications to SAGE data. Biostat. 9, 321–332 (2008).

25. Robinson, M. D. & Oshlack, A. A scaling normalization method for differential expression analysis of RNA-seq data. Genome biology 11, 1 (2010).

26. Robinson, M. D., McCarthy, D. J. & Smyth, G. K. edgeR: a Bioconductor package for differential expression analysis of digital gene expression data. Bioinforma. 26, 139–140 (2010).

27. McCarthy, D. J., Chen, Y. & Smyth, G. K. Differential expression analysis of multifactor RNA-seq experiments with respect to biological variation. Nucleic acids research gks042 (2012).

28. Zhou, X., Lindsay, H. & Robinson, M. D. Robustly detecting differential expression in RNA sequencing data using observation weights. Nucleic acids research 42, e91–e91 (2014).

29. Lun, A. T., Chen, Y. & Smyth, G. K. It’s DE-licious: a recipe for differential expression analyses of RNA-seq experiments using quasi-likelihood methods in edger. Stat. Genomics: Methods Protoc. 391–416 (2016).

30. Anders, S. & Huber, W. Differential expression analysis for sequence count data. Genome Biol. 11, R106 (2010). URL http://genomebiology.com/2010/11/10/R106/. DOI 10.1186/gb-2010-11-10-r106.

31. Bolstad, B. M., Irizarry, R. A., Astrand, M. & Speed, T. P. A comparison of normalization methods for high density oligonucleotide array data based on variance and bias. Bioinforma. 19, 185–193 (2003).

32. Ritchie, M. E. et al. limma powers differential expression analyses for RNA-sequencing and microarray studies. Nucleic Acids Res. 43, e47 (2015).

33. Law, C. W., Chen, Y., Shi, W. & Smyth, G. K. Voom: precision weights unlock linear model analysis tools for RNA-seq read counts. Genome Biol 15, R29 (2014).

34. Liu, R. et al. Why weight? Modelling sample and observational level variability improves power in RNA-seq analyses. Nucleic acids research 43, e97–e97 (2015).

35. Hardcastle, T. J. & Kelly, K. A. baySeq: empirical bayesian methods for identifying differential expression in sequence count data. BMC bioinformatics 11, 422 (2010).

36. Li, J., Witten, D. M., Johnstone, I. M. & Tibshirani, R. Normalization, testing, and false discovery rate estimation for RNA-sequencing data. Biostat. kxr031 (2011).

37. Lund, S. P., Nettleton, D., McCarthy, D. J., Smyth, G. K. et al. Detecting differential expression in RNA-sequence data using quasi-likelihood with shrunken dispersion estimates. Stat Appl Genet. Mol Biol 11, 8 (2012).

38. Hammer, P. et al. mRNA-seq with agnostic splice site discovery for nervous system transcriptomics tested in chronic pain. Genome research 20, 847–860 (2010).

39. Bottomly, D. et al. Evaluating gene expression in C57BL/6J and DBA/2J mouse striatum using RNA-seq and microarrays. PloS one 6, e17820 (2011).

40. Consortium, G. The Genotype-Tissue Expression (GTEx) project. Nat Genet. 45, 580–585 (2013). URL http://dx.doi.org/10.1038/ng.2653. Commentary.

41. Zhang, W. et al. Comparison of RNA-seq and microarray-based models for clinical endpoint prediction. Genome biology 16, 133 (2015).

42. Dillies, M.-A. et al. A comprehensive evaluation of normalization methods for illumina high-throughput RNA sequencing data analysis. Briefings bioinformatics 14, 671–683 (2013).

43. Zyprych-Walczak, J. et al. The impact of normalization methods on RNA-seq data analysis. BioMed research international 2015 (2015).

44. Lin, Y. et al. Comparison of normalization and differential expression analyses using RNA-seq data from 726 individual Drosophila melanogaster. BMC genomics 17, 1 (2016).

45. Subramanian, A. et al. Gene set enrichment analysis: a knowledge-based approach for interpreting genome-wide expression profiles. Proc. Natl. Acad. Sci. 102, 15545–15550 (2005).

46. Xiao, Y. et al. A novel significance score for gene selection and ranking. Bioinforma. 30, 801–807 (2012).

47. Benidt, S. & Nettleton, D. SimSeq: a nonparametric approach to simulation of RNA-sequence datasets. Bioinforma. 31, 2131–2140 (2015).

48. Benjamini, Y. & Hochberg, Y. Controlling the false discovery rate: a practical and powerful approach to multiple testing. J. royal statistical society. Ser. B (Methodological) 289–300 (1995).

49. Bray, N., Pimentel, H., Melsted, P. & Pachter, L. Near-optimal probabilistic RNA-seq quantification. Nat. Biotechnol 34, 525–527 (2016).

50. Edgar, R., Domrachev, M. & Lash, A. E. Gene Expression Omnibus: NCBI gene expression and hybridization array data repository. Nucleic acids research 30, 207–210 (2002). URL https://www.ncbi.nlm.nih.gov/.

51. Soneson, C. & Robinson, M. D. Towards unified quality verification of synthetic count data with countsimqc. Bioinforma. (2017).

52. Chang, W., Cheng, J., Allaire, J., Xie, Y. & McPherson, J. shiny: Web Application Framework for R (2017). URL https://CRAN.R-project.org/package=shiny. R package version 1.0.4.

